# Locating the sex determining region of linkage group 12 of guppy (*Poecilia reticulata*)

**DOI:** 10.1101/2020.04.30.063420

**Authors:** Deborah Charlesworth, Roberta Bergero, Chay Graham, Jim Gardner, Lengxob Yong

## Abstract

We describe new genetic mapping results from 6 full-sib families in the guppy (*Poecilia reticulata*), two of which included recombinants between the X and Y chromosomes. These recombinants confirm that the guppy sex-determining locus is in the region identified by all previous studies, including a recent report suggesting a candidate sex-determining gene in this fish, close to the pseudo-autosomal region (or PAR) at the chromosome terminus. Our results suggest the presence of some errors in the current assembly of the guppy genome. In males, crossing over occurs at a very high rate in the PAR, and our genetic map of the region allows us to correct the marker order. We also identified two unplaced scaffolds carrying genes that map to the PAR. Genetic mapping cannot be used to order markers in the region where crossing over is infrequent. However, our recombinant male is informative about the order, under the reasonable assumption that crossovers are infrequent. Our mapping families and natural population samples also show that the recently proposed candidate for this species’ sex-determining gene is not completely sex-linked. We detect an association between individuals’ sex and an SNP in the sex-determining region, but not with a marker 0.9 Mb away from it, suggesting that variants in this region may be in linkage disequilibrium with the actual sex-determining factor, but that the factor itself has not yet been identified. So far, no consistently male-specific variant has been identified in the guppy sex-determining region.

## Introduction

The sex chromosome pair of the guppy have been studied for many decades with the aim of understanding the evolution of recombination suppression between Y and X chromosomes. The earliest genetic studies in the guppy discovered sex linkage of male coloration traits that are polymorphic in guppy natural populations (Winge 1922b; Winge 1922a), and numerous such traits in natural and captive populations have been shown to be transmitted from fathers to sons in this fish, with only a few coloration factors being autosomal (Haskins and Haskins 1951; Haskins *et al.* 1961; Lindholm and Breden 2002). The number that show complete or (more rarely) partial sex-linkage is disproportionately large, given that this fish has 23 chromosome pairs (Nanda *et al.* 2014), and the sex chromosome pair represents about 3.8% of the genome (Künstner *et al.* 2017), strongly suggesting that these factors are sexually antagonistic. It has therefore been thought likely that they may have been involved in selecting for suppressed recombination between the two members of this chromosome pair.

Cytogenetic studies have shown that the XY pair are not strongly heteromorphic (Winge 1922b), though differences between the Y and X have been detected in some material (Nanda *et al.* 2014). A sex-linked region that includes less than half of the sex chromosome pair has been identified; the region shows heterochromatin, and is stained by male-specific probes in FISH experiments, and the Y sometimes appears longer than the X (Nanda *et al.* 1990; Traut and Winking 2001; Lisachov *et al.* 2015). A study of crossover locations using MLH1 focus detection on chromosomes in sperm cells (Lisachov *et al.* 2015) found localization of crossovers at the termini most distant from the centromeres of the guppy XY pair (and the same general pattern was seen for the autosomes). The chromosomes are acrocentric, and the mean number of MLH1 foci in a testis cell was 23.2 ± 0.5, representing a single crossover on each chromosome on average (Lisachov *et al.* 2015). Genetic mapping with molecular markers also suggested that crossovers on the sex chromosome pair in male meiosis were commoner in the terminal region than in regions nearer the centromere (Tripathi *et al.* 2009). However, the crossover events were not localized physically; nevertheless, dividing the chromosome diagram shown in that paper into 6 intervals, 12 events in males occurred in the terminal interval, and only one in the adjacent interval, versus all 15 events in females being assigned to the 4 more centromere-proximal intervals (this is highly significant by a Fisher’s Exact test, P = 0.0001). (Lisachov *et al.* 2015) also observed a deficit of crossover events in male meiosis, in a terminal part of the XY pair that could be just centromere-proximal to a male-specific heterochromatic region of LG12, where male-specific probe sequences also hybridize (Nanda *et al.* 1990; Lisachov *et al.* 2015). It is therefore likely that this region carries the sex-determining locus (Figure 1).

**Figure 1.**
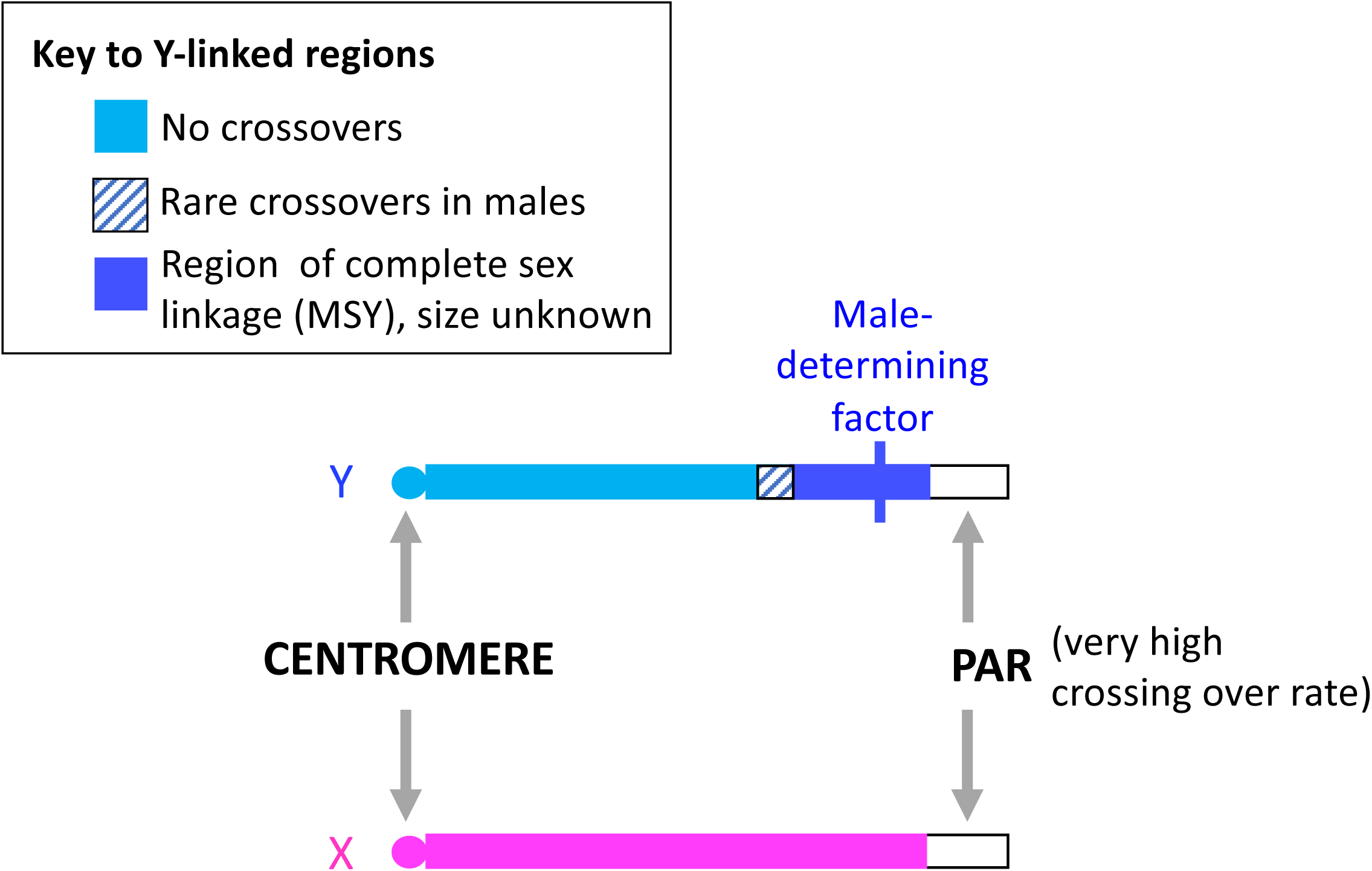
Diagram of the *P. reticulata* XY sex chromosome pair to show regions with different crossover rates in males, based on cytogenetic data and population genomic results (see Introduction). Much or all of the chromosome is present on both the X and Y, and coverage of sequences is very similar in the two sexes (the Y is not degenerated and appears not to have lost genes carried by the X, see (Bergero *et al.* 2019). The diagram shows the centromere at the left-hand end (as in the genome assembly), and the pseudo-autosomal region (or PAR), where most crossover events occur, at the right-hand end; the extent of any pericentromeric region with extremely low crossing over in male meiosis is unknown, but it might extend across much of the XY pair. The Results section below describes evidence that the male-determining locus (symbolized by a blue vertical line) is just proximal to the PAR. It is possible that the male-determining region is missing from the female assembly used in analyses of coverage, and therefore remained undetected (since such male-specific sequences would not have mapped to the female reference genome, including its unplaced scaffolds). It is also not known whether the male-determining gene(s) lie within a completely non-recombining male-specific Y region (or MSY), as the presence of male-specific heterochromatin suggests (see Introduction); if so, the region could also include other genes, as in other species’ MSY regions. It remains possible that there is just a very small maleness factor, for example a single gene or SNP, as discussed in the Introduction section.

However, population genomic studies suggest that variants on chromosome 12 show weak associations with the sex-determining region, although associations are detectable across much, or even the whole, of the XY pair (Wright *et al.* 2017; Bergero *et al.* 2019), and there is evidence for migration of sequences across a large centromere-proximal region between the X and Y chromosomes, strongly suggesting that recombination occurs (Bergero *et al.* 2019); this could reflect rare crossover events in the region proximal to the terminal highly recombining pseudo-autosomal region (or PAR) that was detected cytologically by (Nanda *et al.* 1990; Lisachov *et al.* 2015), see Figure 1. Further family studies are therefore needed to refine our understanding of the guppy sex-determining region. It is currently not clear whether this fish has an extensive, multi-gene fully sex-linked region, or a small region, potentially a single gene like the insertion of the maleness factor in the medaka, *Oryzias latipes* (Kondo *et al.* 2009), or even a single SNP, as has been inferred in another fish, the tiger pufferfish or Fugu, *Takifugu rubripes* (Kamiya *et al.* 2012).

It was recently reported that a candidate for the sex-determining region has been mapped on linkage group 12 of the guppy (Dor *et al.* 2019). Genotypes of a microsatellite marker named *gu1066* within the Stomatin-like 2 (*STOML2*) gene, were found to be associated with the sexes of fish from two domesticated guppy strains, Red Blonde and Flame. This marker is located at 25,311,467 bp in the published assembly of the guppy LG12 (Künstner *et al.* 2017), in the region where the cytogenetic results just outlined suggest the presence of the sex-determining locus. These authors also identified a candidate for the guppy sex-determining gene within a 0.9 Mbp region near this marker. Of 27 annotated genes in his region, the *GADD45G-like* gene (at 24,968,682 bp in the LG12 assembly) was identified as a good candidate for the guppy sex-determining gene, or at least for having with a role in male fertility, although the sequence did not include any male-specific alleles and its expression was not sex-biased (Dor *et al.* 2019).

Here, we show that the *gu1066* microsatellite does not have male-specific alleles in natural populations of Trinidadian guppies, and that a polymorphic SNP (single nucleotide polymorphism) in the candidate gene has no male-specific allele. We also describe new mapping results using multiple microsatellite and SNP genetic markers that narrow down the sex-determining locus to a similar region as that identified by all the previous studies cited above, including Dor et al. (2019).

## Methods

### Fish samples, DNA extraction

Table S1 describes the 14 natural populations samples analysed here, and the sources of the fish used for genetic mapping. These were sampled from natural populations in Trinidad in February 2017, and most individuals were photographed as live specimens in the field before being killed and preserved. Live fish from the same natural populations were also transported to the United Kingdom, and maintained at the University of Exeter, Falmouth, for genetic studies (see below).

Genomic DNA for microsatellite genotyping, genotyping of intron length variants, and for high-throughput genotyping (see below), was extracted using the Echolution Tissue DNA Kit (BioEcho, Germany). Microsatellite markers ascertained from the guppy complete genome sequence assembly were genotyped as described previously (Bergero *et al.* 2019). The primers are listed in Supplementary Table S2, along with the primers for the *gu1066* microsatellite.

### Genetic mapping and high-throughput genotyping

Because guppy females breed best when kept together with other individuals, some of the “families” whose genotypes are reported here were generated from multiple parental individuals, and the parents of five of the six full sibships described below were distinguished using microsatellite markers. The QHPG5 family was generated from two females housed with one male, and the PMLPB2 family from three females and two males; only the LAH family involved a single male and single female.

Using sequences from the guppy female whole genome sequence assembly (Künstner *et al.* 2017), we identified genic sequences found in all 16 Trinidadian guppy individuals sampled from a natural population (10 males and 6 females), and ascertained SNPs across LG12 from our own resequencing of these individuals (Bergero *et al.* 2019). These SNPs were targetted for high-throughput genotyping (SeqSNP) experiments with genomic DNA extracted as described above. The experiments were carried out by LGC Genomics (LGC Genomics GmbH, Ostendstraße 25, 12459 Berlin, Germany, www.lgcgroup.com/genomics). The targetted SNPs were selected from within coding sequences, with the criterion that about 50 bp of sequence flanking each such SNP should also be coding sequence, in order to maximise the chance that the sequence would amplify in diverse populations, and to minimise the representation of SNPs in repetitive sequences. To further avoid repetitive sequences, the SNPs were chosen to avoid ones whose frequencies in the ascertainment sample were 0.5 in both sexes. The SNPs and their locations in the guppy genome assembly are listed in a Supplementary Table S10, together with some features of the mapping results. As expected, the primers worked well for most targeted sequences.

Genetic mapping was done as described previously, using microsatellites ascertained from the guppy complete genome sequence assembly, as well as these SNPs. The set of SNPs included one within the candidate gene, *GADD45G-like*, identified by (Dor *et al.* 2019); the gene is also named LOC103474023, and starts at position 24,968,682 in the guppy LG12 assembly (Künstner *et al.* 2017).

### Analysis of F_ST_ values between the sexes from these populations

The SNPs detected by the SeqSNP experiments were used to estimate F_ST_ values between the sexes. The analysis included all sites with variants within each of the natural population samples, which differed between the samples.

### Analyses to search for potentially sex-linked genes in guppy unplaced scaffolds

In order to try and include the whole of the guppy sex chromosome, we also identified two unplaced scaffolds, NW_007615023.1 and NW_007615031.1, that are located near one end of the assembly of the *Xiphophorus maculatus* (platyfish) chromosome, Xm8, which is homologous to the guppy LG12 (Amores *et al.* 2014). The platyfish is a close relative of the guppy (Pollux *et al.* 2014), with a mean synonymous site divergence of around 10%; for comparison, this is similar to the value for *Drosophila melanogaster* and *D. simulans* (Tamura *et al.* 2004).

To search for such orthologues, we did BLAT searches using as query cDNA sequences of genes on chromosome 8 of the platyfish (the homologue of the guppy sex chromosome pair) against all cds found in the guppy (target), and the reciprocal BLAT search. This identified two unplaced scaffolds included multiple genes (the complete results are shown in the file “Copy of_Unlocalised_Contigs_UNLOCALISED_Scaffolds_LG12_P8_enriched” deposited in Dryad). NW_007615031.1 is about 259 kb, and is assembled at ∼ 5-8 kb from the start of the Xm8 assembly (which corresponds with the non-centromere end of the guppy LG12 assembly), and it includes at least 17 genes, and possibly as many as 34 genes. NW_007615023.1 is a 146 kb scaffold that is assembled 1.4 −1.5 Mb from the Xm8 start, and carries 5-12 genes. We identified and mapped microsatellites within these scaffolds.

## Results

### No complete association with maleness for markers in the GADD45G-like gene region

Table 1 shows the *gu1066* microsatellite genotypes in three natural high predation populations of Trinidadian guppies. 16 individuals (32 alleles) of each sex were sampled from each population, and a total of 29 different alleles were observed, of which 11 were shared between two or more population samples (Tables S3 and S4). None of the alleles showed any association with the sexes of the individuals, and no alleles, other than occasional rare alleles (mostly singletons), were male-specific in any of the populations sampled. No results consistent with complete Y-linkage were seen. For example, allele 264 was seen only in 4 males from the Guanapo population, and one male from the Yarra population, but it is homozygous in one of the Guanapo males, showing that it can be carried on the X as well as the Y chromosome. Because false-positive associations between variants are most likely to be seen in small samples, our samples are sufficient to show that these natural populations do not display any strong association of this region with the sex-determining locus.

**Table 1.** Genotypes at the SNP in the candidate sex-determining gene in the two high-throughput genotyping experiments (the experiment number is indicated in column 1). The river from which each population sample was derived is given, and HP and LP denote high-and low-predation sites, respectively.

| Experiment number | Population | Numbers of individuals |  |  |  |
| --- | --- | --- | --- | --- | --- |
|  |  | Males | Females | Heterozygous males | Heterozygous females |
| 1 | Guanapo LP (GLZ) | 14 | 10 | No variation |  |
| 1 | Marianne HP | 17 | 7 | No variation |  |
| 1 | Marianne LP | 14 | 10 | No variation |  |
| 1 | Yarra HP | 14 | 10 | No variation |  |
| 1 | Yarra LP | 14 | 10 | No variation |  |
| 2 | Aripo HP, AH | 14 | 10 | 1 | 10 |
| 2 | Aripo LP, AL | 14 | 10 | 3 | 10 |
| 2 | QL | 16 | 10 | No variation |  |
| 2 | QH | 16 | 10 | 1 | 10 |
| 2 | TL | 16 | 10 | 6 | 10 |
| 2 | TH | 16 | 10 | 1 | 10 |
| 2 | PL | 8 | 16 | No variation |  |
| 2 | GLT | 0 | 10 | No variation |  |
| Totals across all populations with variants |  |  |  | Heterozygous | Homozygous |
| Total in males |  |  |  | 15 | 75 |
| Total in females |  |  |  | 0 | 60 |

The polymorphic SNP in the candidate gene identified by Dor et al. (2019) is, however, associated with maleness, although no allele is fully male-specific. We genotyped this SNP in 13 natural populations, and it was polymorphic in six of them. Table 1 shows the results from these six populations pooled (the full details, including sample sizes are in Table S5). A Fisher’s exact test shows that, overall, males are heterozygous significantly more often than females (P = 0.0004). However, only 17% of the males genotyped are heterozygotes (Table 1).

### Analysis of F_ST_ values between the sexes in the same natural population samples

14 of the natural populations shown in Figure S1 were genotyped for LG12 SNPs to test whether the SNPs that showed fully sex-linked genotype configurations in a previous study (Bergero *et al.* 2019) do indeed have variants completely linked to the sex-determining locus. Figure 1 shows that, although associations are detected by our analysis, they are not complete; some males are not heterozygous for the variants examined, or some females are heterozygous. Similarly to our previous results using a captive population, F_ST_ values between the sexes are often higher in the centromere-proximal region of the chromosome (proximal to 5 Mb in the published female assembly), and, to a lesser extent, across the region distal to 20 Mb, where the sex-determining locus is thought to be located (see Figure 1). However, as discussed below, genetic mapping and other results show that these signals of associations with the sex-determining locus seen in the proximal region are not evidence that the sex-determining locus is located in a proximal part of LG12.

### Genetic mapping in families to detect the PAR boundary and X-Y recombination events

Out of 721 individuals of known sex that we have genotyped for genetic mapping in twelve *P. reticulata* families (Table S1), only two recombinants were detected, one male in family QHPG5, and one female in family PMLPB2. The dams and sires of both families were sampled alive from natural populations in Trinidad and the progeny were generated in the United Kingdom (see Methods).

The PMLPB2 family (from the Petit Marianne river, in the Northern drainage of the Trinidad Northern mountain range, see Willing *et al.* 2010), and the results of genotyping seven LG12 microsatellite markers are shown in Supplementary Table S6; out of 69 female and 68 male progeny genotyped, the recombinant female (PMLPB2_f23r) carries the sire’s Y-linked alleles at the 4 more proximal LG12 markers, located from 1.2 to 11.7 Mb in the published assembly, but his X-linked alleles at the 3 distal markers, including a marker at 21.3 Mb. This establishes that the sex-determining locus cannot be proximal to 21.3 Mb.

The QHPG5 family dam and sire were sampled from a natural population in the Quare river, in the Atlantic drainage (Willing *et al.* 2010), and the genetic mapping results are shown in Supplementary Table S7. The QHPG5 parents and the small number of progeny (in total, from two full sibships with different dams; sibship 1 included 13 progeny, and sibship 2 had 10 progeny). The genotypes of LG12 microsatellites yielded one recombinant male (male 7, in sibship 2, labelled QHPG5m07r in Table S7) that inherited his father’s X-linked markers for markers spanning the centromere-proximal part of the chromosome, and his father’s Y-linked alleles for markers distal to about 21.2 Mb. To define the crossover breakpoint in male 7, the fish in this sibship (and in the QHPG5 sibship 1) were also genotyped for SNPs throughout LG12, using a high-throughput genotyping approach (see Methods).

Before describing these results, we discuss the PAR boundary, which must be proximal to the sex-determining locus. The PAR boundary was defined in the QHPG5 family based on the most distal gene showing complete co-segregation with sex among all the progeny other than male 7 in both QHPG5 family sibships, an SNP at position 25,998,942 in the LG12 assembly, whose total size is estimated to be 26.44 Mb (Künstner *et al.* 2017). A slightly more centromere-proximal boundary, 25,194,513 bp, is found in two other, larger families, the LAH family previously described, from a high-predation site in the Aripo river, with 42 progeny (see Supplementary Table S1 and Bergero *et al.* 2019), and family ALP2B2, from a low-predation site in the Petit Marianne river, with 136 progeny, see Supplementary Tables S6, S7, S9 and S10).

We next defined the proximal boundary of the crossover event in the QHPG5 family sibship 2, which includes the recombinant male 7. This sibship has the same male parent as sibship 1, but a different dam. Supplementary Figure S7 shows results for both types of markers in both sibships. In sibship 1, 137 LG12 markers informative in male meiosis show complete sex-linkage, while 10 markers from the terminal part of the assembly of this chromosome are partially sex-linked and can definitively be assigned to the small pseudo-autosomal region (or PAR), consistent with previously published results for this family and for seven other guppy families studied (Bergero *et al.* 2019). Figure 4 shows results for both types of markers in sibship 2, which yielded evidence confirming the crossover event suggested by the microsatellite marker results described above. Despite sibship 2’s small size, the crossover is confirmed by twenty markers (16 SNPs and 4 microsatellites), although the data also reveal likely assembly errors, which we discuss in the next section.

**Figure 2.**
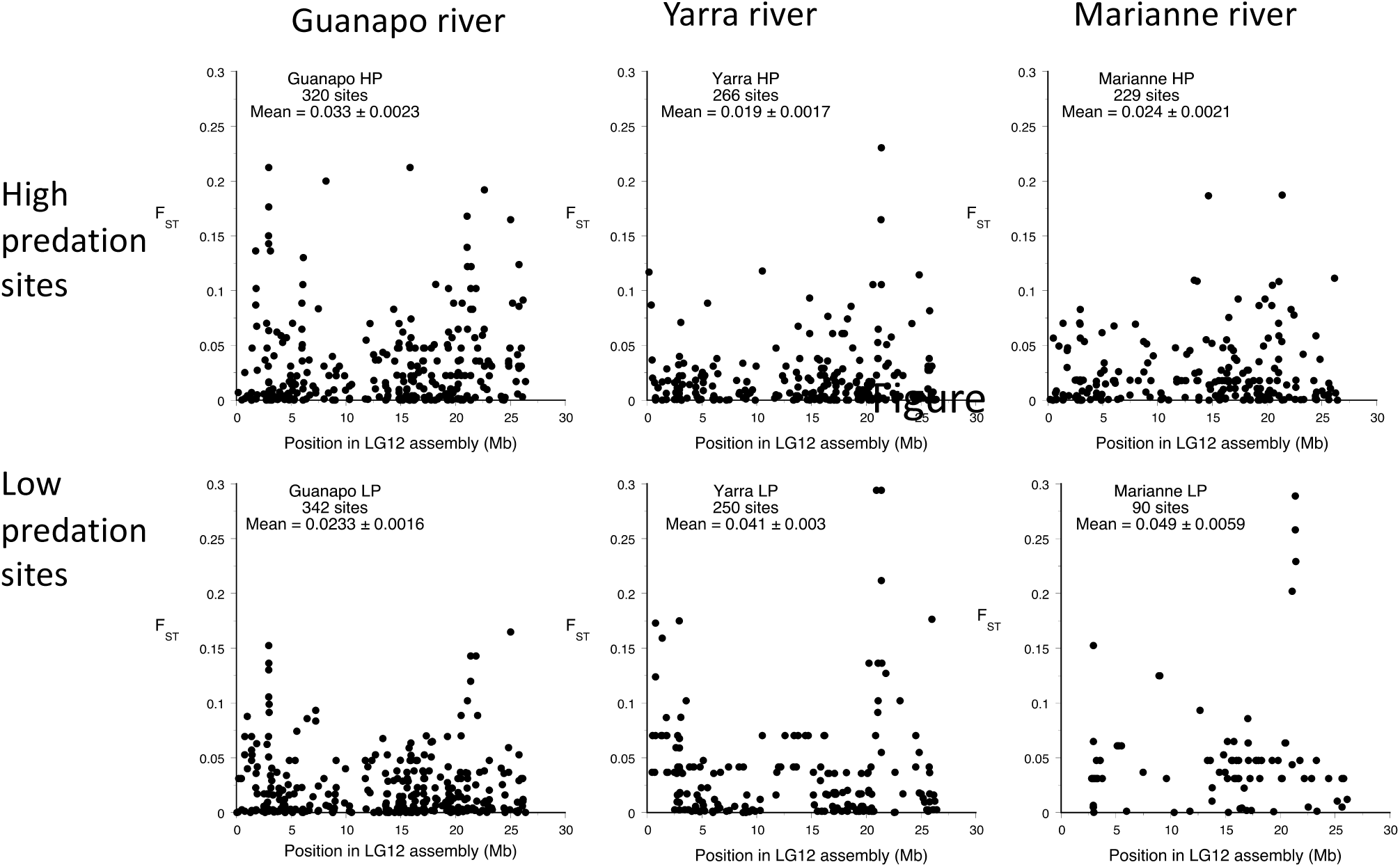
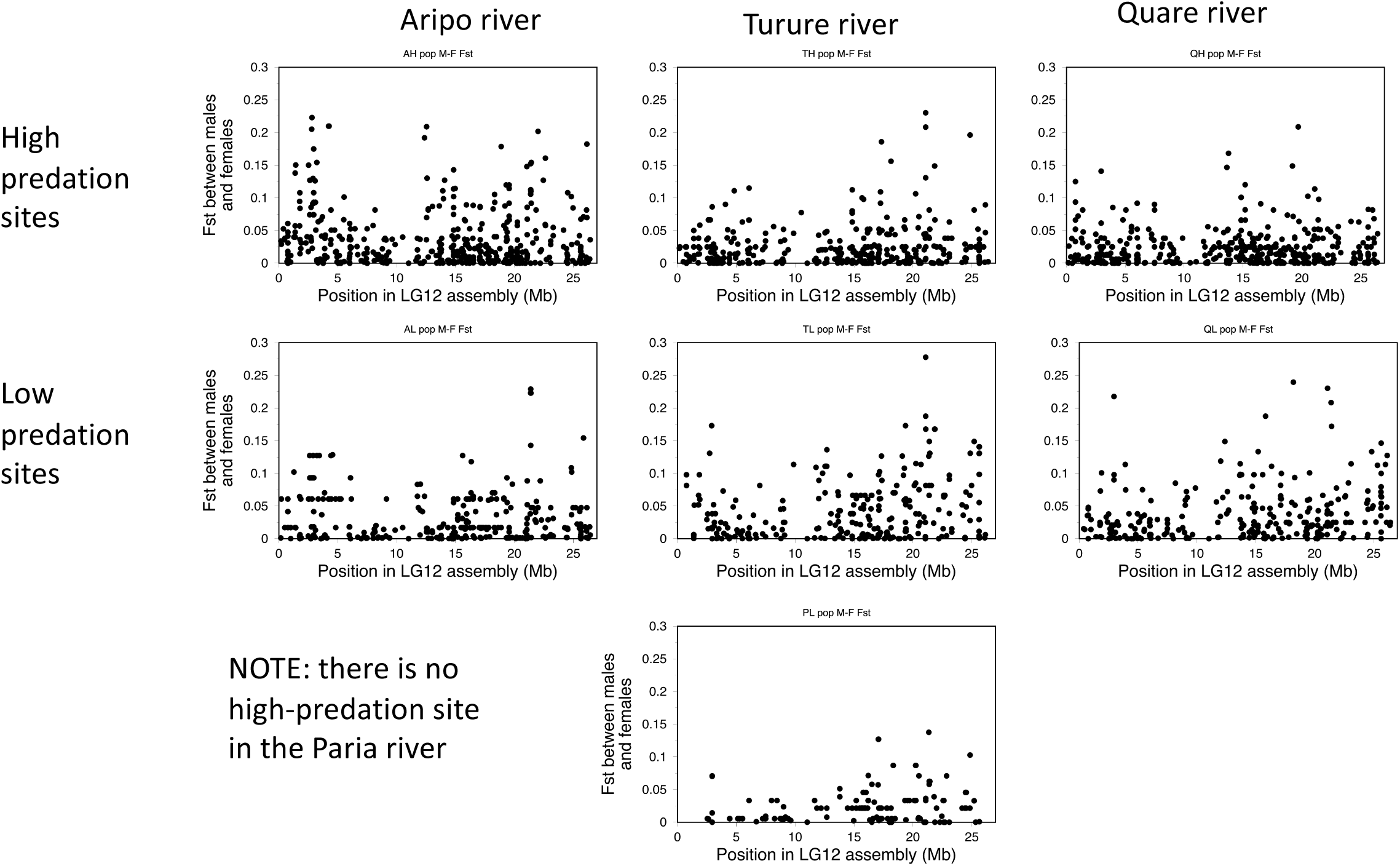
Differentiation between males and females sampled from natural populations from rivers in the Northern range of Trinidad (Supplementary Table S1), based on high-throughput genotyping results for LG12 SNPs.

**Figure 3.**
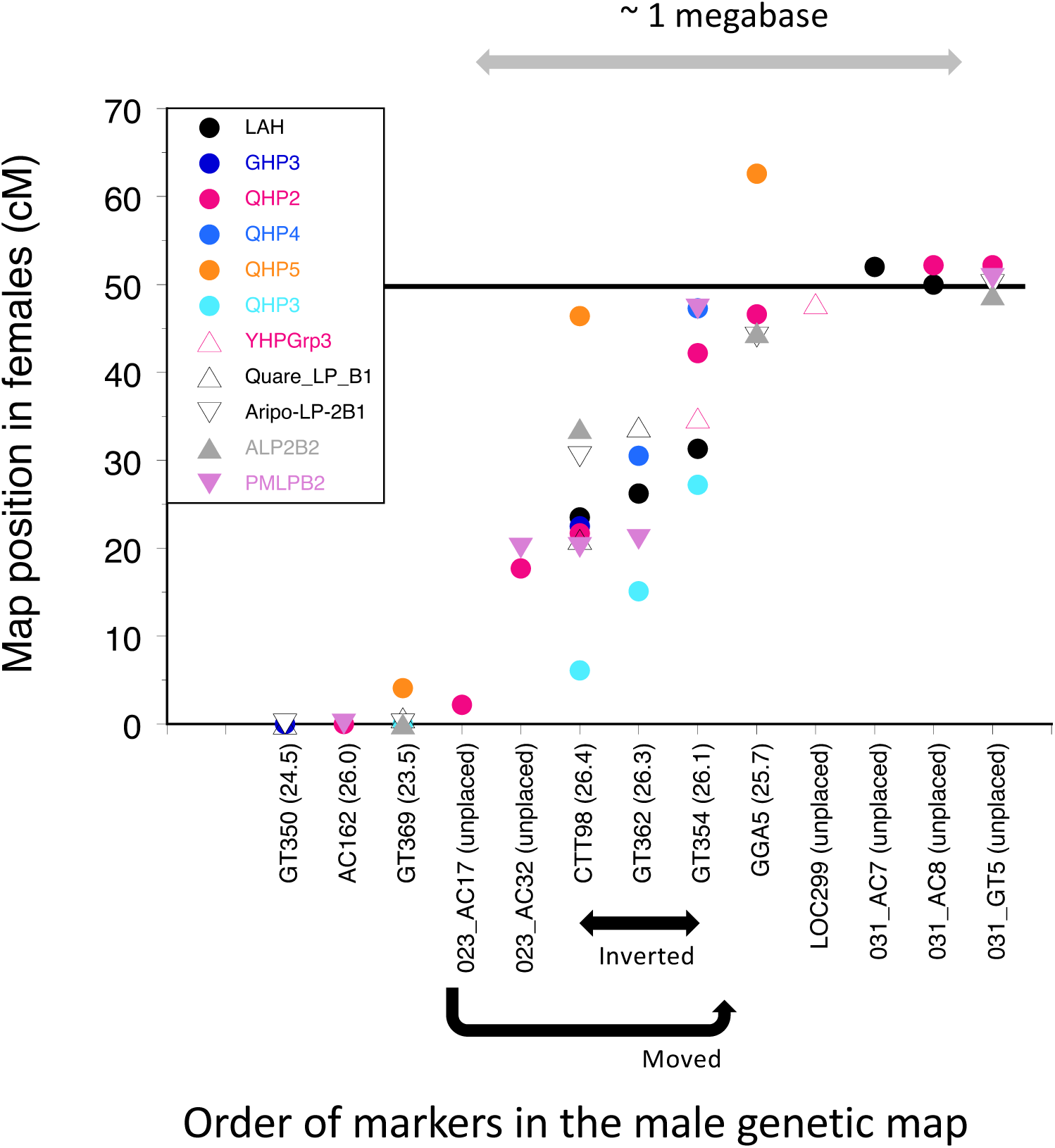
Genetic map of the guppy PAR, based on male meiosis with multiple families with sires from natural populations, in Trinidad (seven from high-predation sites and four from low-predation sites, see Supplementary Table S1). The different colour dots indicate different families. Each family’s dam is from the same population as the sire. Only PAR markers are shown, and all more proximal markers co-segregate with the progeny individuals’ sexes (see Supplementary Figure S2), apart from the single recombinant male in the QHPG5 family (see Figure 4 and the text), and a single female in family PMLPB2 (see text). Note that the marker map order in male meiosis is shown, not the positions of markers on the chromosome in megabases (because the errors in the assembly of this region mean that the true positions are not known).

**Figure 4.**
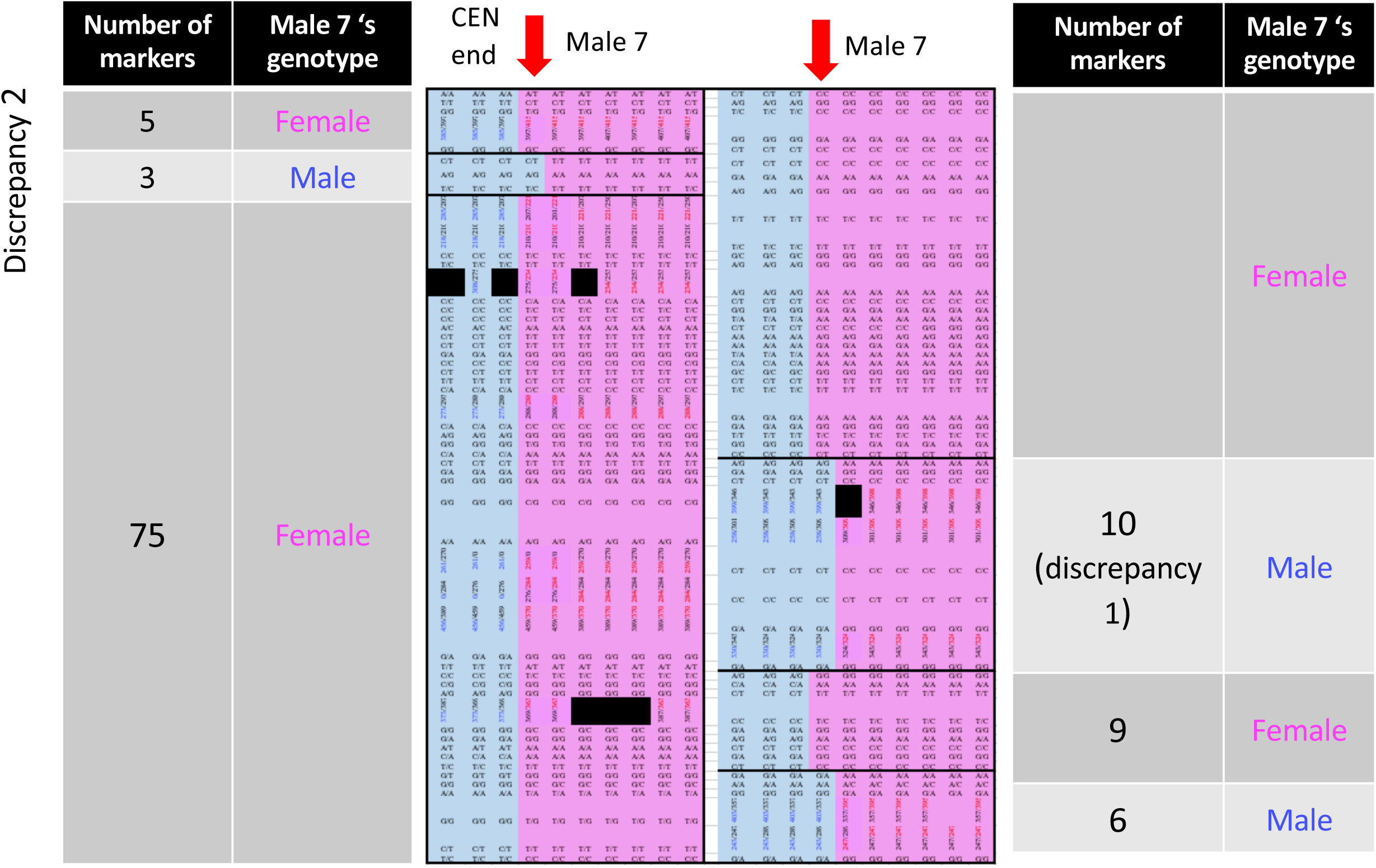
Summary of genotypes at non-PAR markers informative in male meiosis in sibship 2 from the Quare QHPG5 family, in diagrammatic form, showing the two discrepancies between these male genetic map results and the female genome assembly. More details of the same results are given in Table 1. The left-hand part shows the 58 more centromere-proximal markers, and the right-hand part shows the 58 more terminal markers, up to the inferred PAR boundary. The recombinant male (male 07) is indicated at the top of the diagram in both parts, while the other columns in both parts show genotypes at the 108 markers that, in the 9 other progeny, co-segregate perfectly with the individuals’ sexes in meiosis of the male parent (no recombinants were found in the 13 sibship 1 progeny, which have the same sire, as shown in Supplementary Table S7).

In sibship 2, 141 markers appear to be sex-linked, with all female progeny inheriting the same paternal allele, which is therefore the sire’s X-linked allele, and three males all inheriting the other, Y-linked, paternal allele. However male number 7 is recombinant: he inherited the paternal X-linked allele for the first 111 markers (with 12 exceptions, discussed below), but the paternal Y-linked allele for 21 markers informative in male meiosis, 14 in positions from 21,049,596 to 23,293,233, and seven from 24,829,827 to 25,998,942 bp (after which the 10 terminal markers appear to be partially sex-linked, as in the other families). The male-determining region must therefore be distal to position 21,049,596 in the assembly, and centromere-proximal to the position of the first PAR marker. We next discuss information from which we can infer the correct order of some of the markers, and determine which PAR marker is the most proximal of those so far mapped.

### Detection and correction of assembly errors in LG12: the PAR

Crossing-over occurs at a high rate in the physically small guppy PAR (Bergero *et al.* 2019), allowing us to order the markers by genetic mapping of the region. We also mapped microsatellites within two unplaced scaffolds that are located near one end of the assembly of the homologous chromosome in the closely related fish, *X. maculatus* (see Methods), revealing that they are part of the guppy PAR. The most proximal PAR marker so far mapped is in the unplaced scaffold, NW_007615023.1, while the NW_007615031.1 scaffold maps terminal to the microsatellite markers mapped previously (Bergero *et al.* 2019).

We were able to order the microsatellite PAR markers because some of them were mapped in several different families, and showed consistent ordering that differed from the guppy assembly, suggesting that this region includes several assembly errors (Figure 3). These errors make it impossible to relate the genetic to the physical map, and therefore the figure simply orders the markers according to their genetic map positions. Based on the total size in the published assembly, plus the two unplaced scaffolds that we infer from the *X. maculatus* assembly belong to the guppy PAR (which add about 405 kb), our results do not change the conclusion that the guppy PAR is not larger than a couple of megabases, and probably smaller (Bergero *et al.* 2019). Assuming that consistent high intronic high GC content of a region reflects a high recombination rate that causes GC-biased gene conversion (see accompanying manuscript), the uniformly high GC content (Supplementary Figure S1) of the scaffold that our map (Figure 3) reveals to be terminal suggests that recombination rates are uniformly high throughout the guppy PAR, rather than being clustered into parts of the PAR with very high crossover rates.

### Correction of assembly errors in the non-PAR part of LG12

Genetic mapping cannot order markers in genome regions where crossovers do not occur, or occur very rarely, as is the case in guppy male meiosis for the LG12 centromere-proximal region of at least 24 Mb of that almost completely co-segregates with the sex-determining locus; Supplementary Figure S2 shows the results for the families reported previously (Bergero *et al.* 2019), plus some others newly mapped here, whose source populations are listed in Supplementary Table S1. However, if crossovers are infrequent, then errors in the ordering of markers can be detected in rare recombinant individuals, because co-segregation of such markers with others can reveal their true physical locations. Our sibship that includes the recombinant male suggests several discrepancies in the order of sequences within the proximal region of at least 24 Mb of the guppy LG12 that largely co-segregates with the sex-determining locus (see Supplementary Table S7).

The region just proximal to the PAR (Figure 3, Table 2 and Supplementary Figure S7), includes two sets of markers in the recombinant male suggesting assembly errors. First, nine SNP markers informative in the sire of this sibship with LG12 assembly positions between 24,166,053 and 24,508,654 bp (a span of 342,601 bp, including SNPs in four separate genes mapped in this sibship) clearly appear to belong more proximally than the surrounding markers, as their segregation patterns in the recombinant male are identical to those for 90 of the 93 more proximal informative markers (Discrepancy 1 in Table 2). The only exceptions to this pattern are three SNPs at positions between 2,952,176 and 2,953,254 (Discrepancy 2 in Table 2), which were included in our genotyping experiment because their genotypes in a set of 10 males and 6 females whose complete genomes we sequenced (see Methods) suggested complete sex-linkage (Bergero *et al.* 2019). Their genotypes in the recombinant male clearly indicate that they are located terminal to the recombination breakpoint that is supported by the 20 markers (16 SNPs and 4 microsatellites mentioned above), where the recombinant male inherited the dam’s X-linked alleles. They cannot be located more precisely, because markers in this region co-segregate in this small sibship.

**Table 2.**
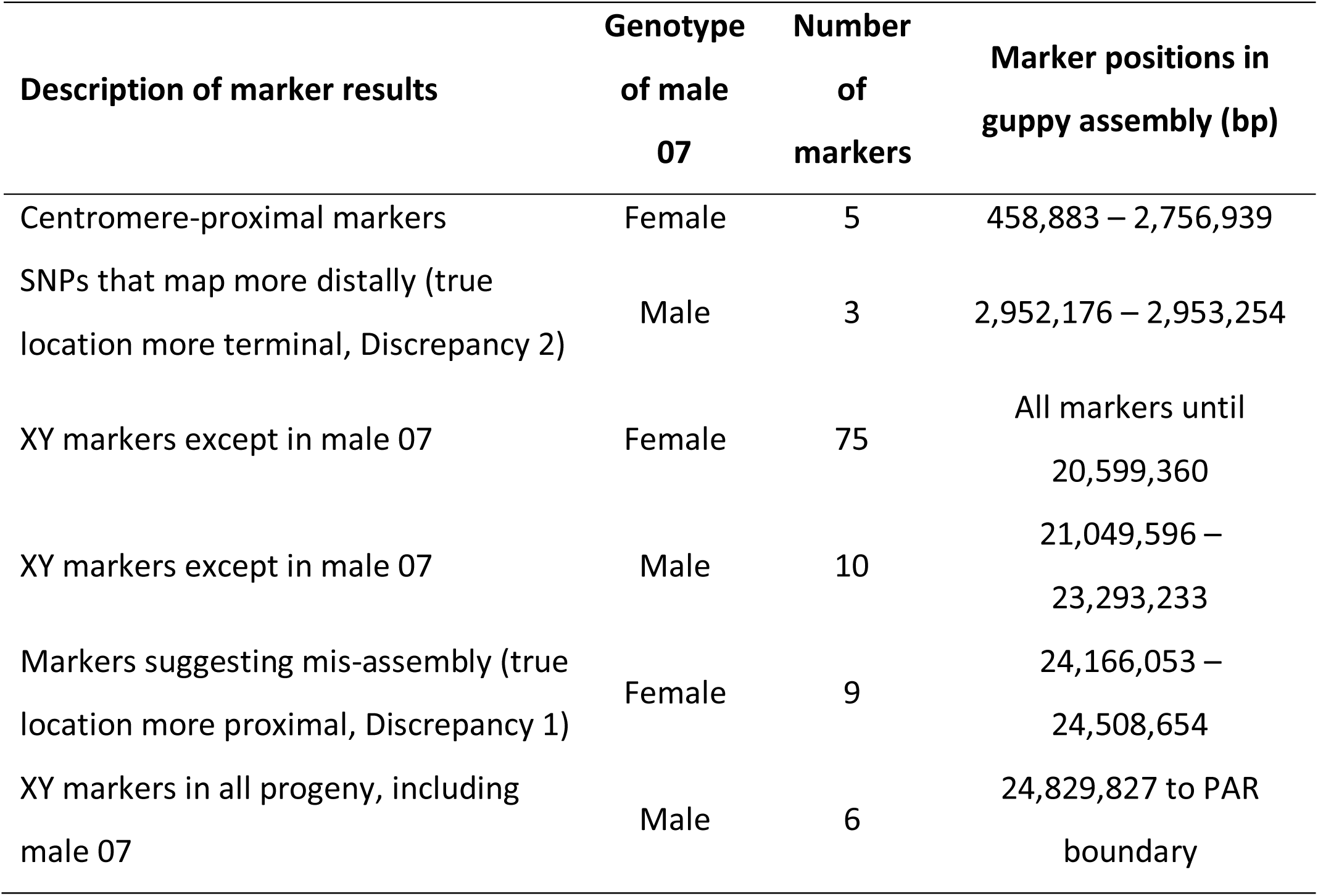
Summary of genotypes in the QHPG5 sibship 2, which includes a recombinant male, for markers that are informative in male meiosis. The two discrepancies are discussed in the text, and the complete genotype information for these markers in both QHPG5 sibships is in Supplementary Table S7. The table shows only markers that co-segregated with the phenotypic sex in all progeny other than male 07, and in all other families where the genotypes were informative in male meiosis. When the recombinant male inherited his sire’s Y-linked allele, his genotype is listed as male in the second column, and it is listed as female when he inherited his sire’s X-linked allele.

To further understand the gene order differences between the high-throughput genetic map results for the QHPG5 family and the guppy LG12 assembly, we searched in the assembly of the homologous platyfish chromosome (chromosome 8) to find the positions of the four genes with the 9 SNPs that constitute Discrepancy 1 in Table 2 and Figure 4. This reveals that two of these are within a region that is assigned a more proximal location in the platyfish assembly (near 15 Mp), near the breakpoint (at 15,847,876 bp) of a region whose order in the genetic maps of all guppy families so far studied that have markers in the relevant region is inverted relative to the published assembly (this includes a map of the LAH family based on high-throughput genotyping, see Supplementary Figure S2). We conclude that the genetic map arrangement based on the guppy QHPG5 family (with these SNPs more proximal than in the guppy assembly) is probably correct, and that these genes should be re-assigned to a more proximal location. The comparison with the platyfish chromosome 8 assembly also indicates that a set of guppy genes that is assembled in megabase 10-11 correspond to ones that are at the terminus of the platyfish chromosome. The AC162 microsatellite marker was based on sequence in this region, but its genetic map location is around 30 cM in female meiosis in the ALP2B2 family, consistent with a location more distal than 11 Mb; this marker has so far been mapped only in this family.

Our families also support the suggestion in Table 2 that the markers in the LG12 region near 3 Mb labelled Discrepancy 2 are located more distally, as the segregation patterns of SNPs that are heterozygous in female meiosis in the dams of three sibships match those of distal markers in the families (SNPs with similar segregation patterns to those of these three SNPs are all distal to position 20,516,823 bp in the LAH family, to 22,002,472 bp in two informative ALP2-B2 family sibships, and to 21,344,716 bp in the QHPG5 family sibship 1). The segregation pattern observed is illustrated in Table 3, together with our interpretation that there is a duplication, which can explain why all males appear to be heterozygous. Furthermore, the segregation patterns of these SNPs in female meiosis in three sibships whose female parents are also heterozygous show that the variants genotyped are located among markers that are close to the PAR boundary, very distant from their assembly positions (Supplementary Tables S8 and S9, for the LAH family and the two informative ALP2B2 sibships, respectively); the same can be seen in the smaller QHPG5 family sibship1 (Supplementary Table S7).

**Table 3.** Example from the LAH family of segregation results found for all three SNPs named “Discrepancy 2” (see text and Table 2). The LAH family sire is inferred to be heterozygous for all three markers. The example shown is for the LG12 site at 2,953,218 bp. The same pattern was observed whenever both the sire and dam had heterozygous genotypes

| Dam<br>genotype | Female progeny |  |  | Sire<br>genotype | Male progeny |  |  |
| --- | --- | --- | --- | --- | --- | --- | --- |
| A/G | A/A | A/G | G/G | A/G | A/A | A/G | G/G |
| Observed<br>numbers | 6 | 11 | 0 |  | 0 | 25 | 0 |
| Interpretation |  |  |  |  |  |  |  |
| $X^A/X^G$ | $X^A/X^A$ | $X^A/X^G$ | $X^G/X^G$ | $X^A/Y^{GY^A}$ | Not<br>expected | $X^A/Y^{GY^A}$<br>$X^G/Y^{GY^A}$ | Not<br>expected |

## Discussion and Conclusions

We conclude from our analysis of genotypes in natural guppy populations that the genetic associations detected (Dor *et al.* 2019) probably reflect linkage disequilibrium within a genome region that recombines rarely with the sex-determining locus, similar to the evidence for incomplete associations of variants with the sexes of individuals recently reported for many parts of the guppy sex chromosome pair (Wright *et al.* 2017; Bergero *et al.* 2019). Dor et al. (2019) concluded that the guppy sex-determining region is a region of 1.26 Mb that is duplicated between LG 9 and 12 and includes 59 LG12 genes, 17 of which had multiple copies; 8 of these had copies on LG 9 as well as 12, while 9 had multiple copies on LG12. The duplicated region occupies 0.43 Mb at about 25 Mb on LG12, and 6 of the 11 genes in the region have multiple copies. We too found a marker that had been assigned to LG9 (the microsatellite GT443) but behaves as fully sex-linked in the male parent of one of our mapping families, LAH (Bergero *et al.* 2019). Its LG9 location is 8.0 Mb, not close to the LG9 region at 17 Mb identified by Dor et al. (2019). Thus the LG9 copy could represent mis-annotation rather than the duplication Dor et al. (2019) identified.

Genes that have moved onto the guppy LG12 since the divergence of the guppy from the platyfish should be found on a platyfish chromosome other than Xm8, the homologue of the guppy LG12. We therefore examined genes on the platyfish chromosome homologous to the guppy LG9 (see Methods). These genes are mostly carried on Xm 12, with a few on chromosome 11 (Supplementary Figure S3A). We inspected the plots for these chromosomes, to determine which guppy chromosomes carry them, and whether some of them are LG12 genes in the guppy. Some platyfish chromosome 11 genes are detected on guppy LG9, but most of them are on LG14 (Supplementary Figure S3C). Platyfish chromosome 12 genes are therefore most relevant for asking whether they form a duplicated region on guppy LG12. Genes on this platyfish chromosome are (as expected) mostly on guppy LG9, with a few on LG25, but a cluster of them are indeed detected on LG12 (Supplementary Figure S2). These genes could well be located in the terminal part of the guppy LG12, which might correspond to the region identified by Dor et al. (2019). The gene order in the centromere-proximal 20 Mb of the 26.4 Mb guppy LG12 is very similar to that in the homologous platyfish chromosome (Xm8, see Supplementary Figure S4). The large inverted region is an assembly error, as the guppy genetic map supports a gene order in agreement with the platyfish assembly (Supplementary Figure S2). Therefore, the location for this set of genes must be terminal to the repeat region just after 20 Mb that is prominently visible in Supplementary Figure S4. The figure also shows that the LG12 assembly terminal to this region is either rearranged, relative to the platyfish homologue, or, in parts, remains uncertain.

Dor et al. (2019) also suggested the “growth arrest and DNA damage inducible gamma-like” (*GADD45G-like*) as a plausible “candidate gene for its role in male fertility”, despite finding “no sex difference in either the genomic sequence or gene expression”. However, several of their observations are difficult to reconcile with this conclusion. First, they state that “the male-specific allele was identified in only 85% of males, but not in females”, and that family D suggested a possible environmental effect of elevated temperature leading to sex reversal, or recombination that produces inviable recombinant females. Furthermore, the *gu1066* marker identified 97 females as genetically XY, suggesting that sex reversals occur; when mated with normal XY males, these should yield 25% females, but only 19% of the progeny had ovaries, and only four live fingerlings were produced (by one sex reversed female), three XY and one XX.

Our results also do not support the conclusion that this is the guppy sex-determining gene, or maleness factor, but only that an SNP in the gene has a variant in a few males from several natural populations that is not seen among females. SNPs in this gene show sex-linkage in the QHPG5 and ALP2B2 families, but none of the our LG9 markers that are informative in male meiosis does (data not shown). These results do not definitively rule out the possibility that the LG12 *GADD45G-like* gene is the guppy male-determining factor. The guppy might have a single SNP that determines maleness, as has been inferred in fugu; within the gene with the mis-sense mutation that apparently controls maleness in fugu (anti-Müllerian hormone receptor type II, or *Amhr2*), only 5 SNPs other than the mutation itself showed (incomplete) associations with individuals’ sexes (all within the surrounding region from 7.27 to < 10 kb, see Figure 2C of (Kamiya *et al.* 2012).

Our new results do, however, support all previous results, including those of from the captive material studied by Dor et al. (2019), suggesting that the guppy male-determining factor is located within a region between about 21 and 25 Mb (in the current assembly). We describe the first genetic data based on localising X-Y crossovers within families, and the two recombinant individuals both indicate that the male-determining factor is located distal to 21 Mb, and our results also define the PAR boundary, which our families indicate is at about 25 Mb (though assembly errors make it difficult to define its boundary or size precisely). Given these problems, and the rarity of recombinant progeny, it remains difficult to define the male-determining locus more accurately. If a more reliable assembly can be generated by long-read sequencing, the rarity of recombination events may still make it impossible to define the locus genetically, though this approach might narrow the region down enough to suggest candidate genes for further testing. Unfortunately, although the *GADD45G-like* gene appears a promising candidate, several observations make it seem unlikely to be the true guppy sex-determining gene.

## Acknowledgements

This project was supported by ERC grant number 695225 (GUPPYSEX). We thank Bonnie Fraser (University of Exeter) for discussions about the large inversion on the guppy LG12.

## Supplementary files

### Supplementary figures

Figure S1. GC content of introns in two scaffolds that were unplaced in the published guppy genome assembly, but which map to the PAR, as shown in Figure 2 of the main text.

Figure S2. Genetic maps of the guppy LG12 in female (left) and male (right) meiosis, in the same families as shown in Figure 3 of the main text.

Figure S3. Comparison of the gene contents of guppy and platyfish chromosomes.

Figure S4. Comparison of the gene arrangements of the guppy LG12 and the homologous chromosome of the platyfish, Xm8.

### Supplementary tables

Table S1. Trinidadian guppy natural population samples studied, including live fish transported to the United Kingdom, and maintained at the University of Exeter, Falmouth, for genetic studies, together with the names of families used for genetic maps. Hgh-throughput experiment 1 included the LAH family. Where possible, both high and low predation populations were sampled from each river; the Paria river has no high predation sites.

Table S2. Primers for the guppy LG12 markers genotyped (mostly microsatellites), including the gu1066 microsatellite and those identified from Xm8, NW_007615031.1 and NW_007615023.1 (see main text).

Table S3. Summary of gu1066 genotypes in three natural populations. Blue indicates males and red females.

Table S4. Table S4. Genotypes found in 32 alleles sampled from each of three high predation populations (16 from each sex). Grey shading indicates alleles with > 3 copies in a population sample, and bold font indicates alleles that were found in more than a single population.

Table S5. Results of high-throughput genotyping for the SNP at site 24,969,110 in the LOC103474023 gene identified by Dor et al. (2019) as a candidate for the guppy sex-determining gene. HP indicates locations with high predation rates and LP indicates low predation rate.

Table S6. Genotypes of seven LG12 microsatellite markers proximal to the pseudo-autosomal region (PAR) in the PMLPB2 family, showing an X-Y crossover event in female progeny PMLPB2_f23. For clarity, PAR markers are not shown. 137 progeny were genotyped, excluding two contaminants whose genotypes do not correspond to any othe sires or dams. The dams and sires of the progeny were inferred from the genotypes, and are shown in columns C and D. The positions in the guppy genome assembly are shown below each marker name. The 68 male progeny (blue font in column A) all carry the variant that can be inferred to be the sire’s Y-linked allele, and 68 of the female progeny inherited their sire’s X-linked allele; two males and one female disagreed with heir assigned sex for all seven markers, and were probably labelled erroneously). Female f23, however, carries her sires Y-linked alleles at the 4 markers from 1.2 to11.7 Mb, and his X-linked alleles at the 3 distal markers, including the marker at 21.3 Mb.

Table S7. Genotypes of 264 LG12 markers in the QGPG5 family, which includes an X-Y crossover event in male progeny QHPG5m07 (in sibship 2). Most of the markers are SNPs, which were genotyped by high-throughput experiments (see main text), but 17 microsatellites were also genotyped. The two sibships had the same sire, but different dams, and the progeny were assigned to their respective dams using the microsatellites. Column B shows the positions in the published guppy genome assembly (with the microsatellite names and positions in bold font). Column A notes several “landmarks”, including locations and sets of markers that are described in the main text. All 13 progeny in sibship 1 show complete co-segregation in male meiosis of all informative markers in the region centromere-proximal to the PAR boundary; marker genotypes are coloured in blue or pink, to indicate whether they inherited the sires’s Y- or X-linked alleles, respectively. Cells coloured in grey indicate SNPs that are not informative in a sibship (markers that were uninformative in both male and female meiosis in both sibships are not shown), and yellow indicates discrepancies from Mendelian expectations, probably due to genotyping errors; these are very infrequent. In sibship 2, one individual (indicated in yellow) carries the sire’s X-linked markers, and is therefore inferred to be a female, although it was recorded as a male, but three males and five other females show complete co-segregation of all non-PAR markers with sex. Male m07, however, is clearly recombinant: this male carries his sire’s X-linked alleles at most centromere-proximal markers, but Y-linked alleles at markers distal to 21.2 Mb, with few exceptions which are discussed in the main text (see Figure 3 for a summary of this sibship’s results for markers informative in male meiosis only).

Table S8. LAH family results from high-throughput genotyping. Colouring is the same as described for Table S7. As there are only a single sire and single dam for this family, all SNPs that are not informative have been omitted. The three SNPs that are boxed (at positions 2,952,176, 2,953,218, and 2,953,254) are discussed in the text under the term “Discrepancy 2”,, and the most proximal position that is consistent with their segregation patterns in female meiosis is distal. to 21 Mb in the assembly. For these three discrepant SNPs, the segregation in male meiosis is not informative about their location, because there is a duplication such that all male progeny of female heterozygotes appear to be heterozgous, as explained in the main text.

Table S9. Segregation patterns in two ALP2B2 sibships in which the three boxed SNPs (at positions 2,952,176, 2,953,218, and 2,953,254) named “Discrepancy 2” are heterozygous in the dams. These are discussed in the text under the term “Discrepancy 2”, and again the most proximal position that is consistent with their segregation patterns in female meiosis is distal. The colouring is the same as described for Table S7. The table shows that, while the segregation patterns for most SNPs in female meiosis agree with those assembled in physically nearby positions (for sibship 1, agreement is seen for all 30 SNPs proximal to the 3 discrepant ones, and for 7 more distal SNPs before the first detectable crossover), this is not the case for these three SNPs. Instead, in both these sibships (totalling 33 female progeny genotyped), their segregation patterns agree closely with those of SNPs at positions near 23 Mb. For these three discrepant SNPs, the segregation in male meiosis is not informative about their location, because there is a duplication such that all male progeny of female heterozygotes appear to be heterozygous, as explained in the main text.

Table S10. Genes on the guppy LG12 (the *Poecilia reticulata* sex chromosome, and their locations in the published guppy genome assembly and in the assembly of the homologous chromosome (Xm8) of the platyfish, *Xiphophorus maculatus*, together with some results from genetic mapping using SNPs selected from across the guppy assembly. The homologous genes in the two species were determined by NUCmer analysis (Kurtz et al., 2004, http://mummer.sourceforge.net) using the platyfish chromosome 8 as the reference and the guppy LG12 sequence as the query. The genes are shown in the order in the published assembly for the guppy, with the guppy centromere end (determined from genetic mapping in male meiosis) at the end with the lowest positions; note that the homologous chromosome if the platyfish is assembled in the opposite order, and therefore distances from the centromere end are shown (in column C), in addition to the assembly positions. Breaks in the platyfish ordering are indicated by grey shading of the regions. 608 sequences with % identity < 85% were excluded. A few SNPs that were mapped in the guppy, but where the locus was not identified in the platyfish, are indicated by turquoise blue text in columns E and F. The large region in which the guppy order is inverted is boxed (as explained in the text, genetic mapping in several families indicates that this is an error in the guppy assembly, rather than a difference between the two species). The PAR is also boxed (based on results in all families with map information), as is the region named “Discrepancy 1” in the main text, and the corresponding positions in platyfish assembly are shown in red, as are the informative guppy SNPs. The region in which the recombinant male in the QHPG5 sibship carries paternal Y-linked variants is highlighted in blue in columns D and E (in all other regions, including the “Discrepancy 1” region, this male carries maternal LG12 alleles, as described in the text).

